# Analysis of Inbred Mouse strains’ High-Impact Genotype-phenotype Hypotheses (AIMHIGH) reveals novel disease-causing candidate genes

**DOI:** 10.1101/2022.08.07.503105

**Authors:** Boyoung Yoo, Surag Nair, Zhuoqing Fang, Rushil Arora, Meiyue Wang, Gary Peltz, Gill Bejerano

## Abstract

Inbred mouse strains reveal the molecular basis of mammalian traits and diseases, particularly recessive ones. We utilized mouse community curated resources to set up an automated screen to discover novel testable gene function hypotheses. Using 11,832 community contributed strain-differentiating experiments and trait presence/absence scoring, we searched for all experiments where strains can be split by their phenotypic values (e.g., high vs. low responders). Then, using 48 sequenced strains, we found one or more candidate gene for each experiment where homozygous high-impact variants (such as stopgain, frameshifts) segregate strains into these same binary grouping. Our approach rediscovered 212 known gene-phenotype relationships, almost always highlighting potentially novel causal variants, as well as thousands of gene function hypotheses. To help find the most exciting hypotheses, we improved the state of the art in machine learning driven literature-based discovery (LBD). Reading on our top 3 ranked candidate genes per experiment reveals 80% of rediscovered relationships, compared to 5% reading at random. We proposed 1,842 novel gene-phenotype testable hypotheses using our approach. We built a web portal at aimhigh.stanford.edu to allow researchers to view all our testable hypotheses in detail. Our open-source code can be rerun as more sequenced strains and phenotyping experiments become available.

## Introduction

Mouse is one of the most widely used model organisms in genetics. Countless biomedical discoveries have been made initially with the mouse model^1^. Inbred laboratory mouse strains are populations of mice that have reached intra-strain genetic homogeneity from at least 20 generations of sibling mating^2,3^. Through generations of inbreeding, each strain has developed unique variants and phenotypes. These carefully maintained populations of mice are expected to be homozygous in most genetic loci, helping expose many recessive inherited traits.

The mouse community has been accumulating a wide range of functional information about mouse genes gained from mouse model experiments. Mouse Genome Database^4^ (MGD) has been at the forefront of curating and organizing this data. They developed Mammalian Phenotype (MP) ontology^5^, which is a structured dictionary of terms that represent various phenotypes observed in mice. Having a controlled set of terms allows for easy and consistent comparisons across annotations.

Inbred mouse strains are often phenotypically and genotypically diverse^6,7^, and many experiments have been performed on these strains to measure their phenotypic diversity. Mouse Phenome Database^8^ (MPD) has curated and publicly released the results of thousands of such experiments deposited from various laboratories covering many different inbred strains.

In addition, MGD has annotated naturally occurring phenotypes in multiple strains. These phenotypes vary widely, ranging from quantitative phenotypes such as time spent in a maze to categorical phenotypes such as coat color.

Here, we analyzed whole genome sequencing data of 48 mouse inbred strains in search of high-impact homozygous variants that segregate concordantly to each phenotypic measure. We then used annotations of causative genes established through knockout experiments to find previously known proof of principle gene-phenotype relationships in our set. To help propose the most exciting hypotheses from a set of novel candidates, we built a literature-based discovery (LBD)^9,10^ classifier trained to predict future single-gene mouse knockout results from clues in the current literature. Our entire pipeline is outlined in Figure 1. Finally, we offer an easy-to-use web interface to display full details of our hypotheses at: aimhigh.stanford.edu.

**Figure 1.**
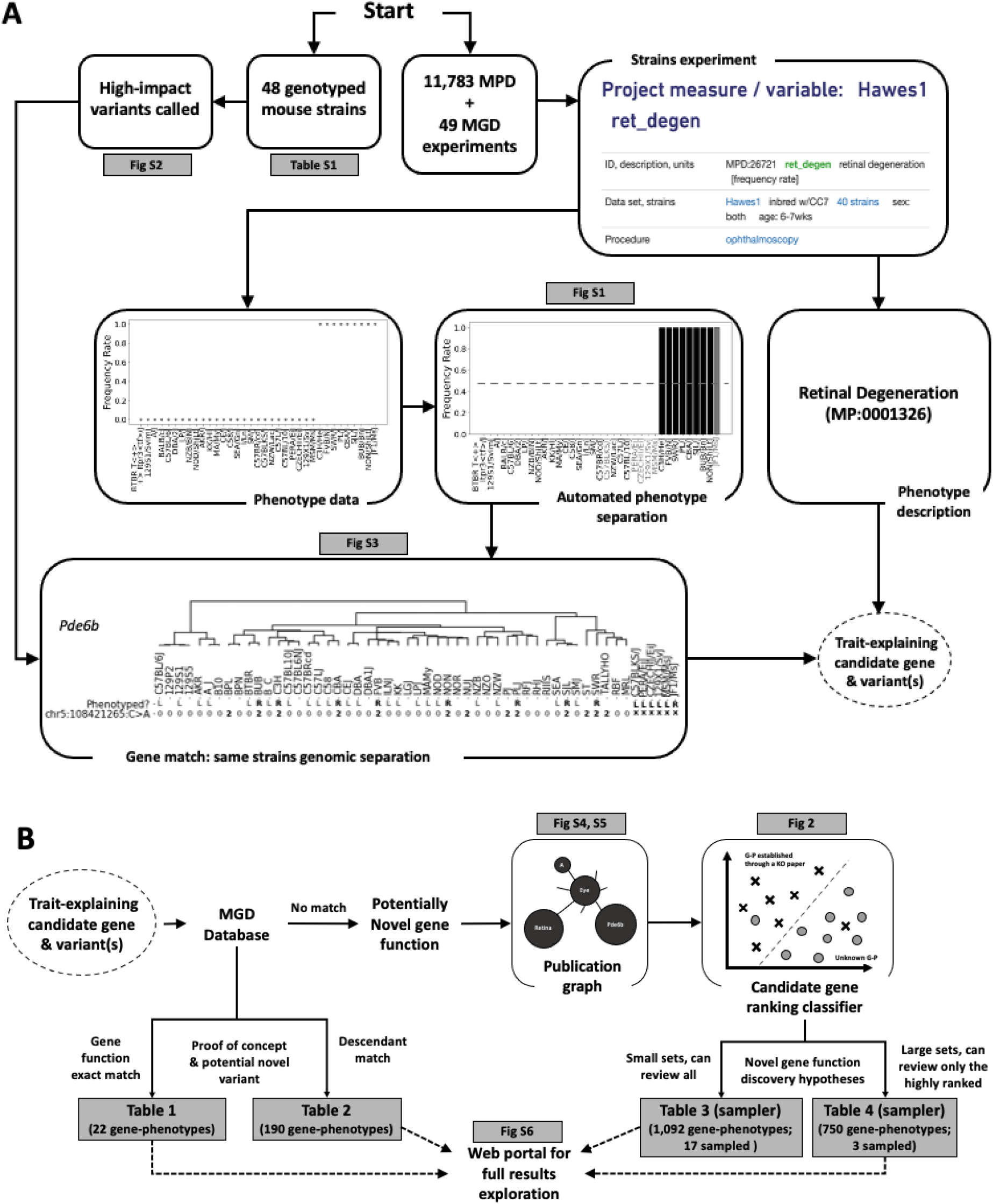
Paper overview. Grey boxes (e.g., Table S1,Fig 2) indicate illustrations and results accompanying each step. **(A)** For every experiment with multi-strain data in Mouse Phenome Database (MPD) and strain-specific presence/absence phenotypes annotated by Mouse Genome Database (MGD), we automatically sought trait values that would partition all measured strains into two groups. For each such phenotypic split, we sought one or more mouse gene with a matching genotypic split, using only the high-impact genomic variants that likely severely modify protein products. Each such match constitutes a trait explaining, possibly novel, hypothesis made by our approach. **(B)** Next, we used MGD known gene function annotations to find hundreds of gene function hypotheses we made that have already been successfully validated by single gene knockout experiments. In all but two such cases, even when the gene level function was already known, the high-impact variant(s) we discover in different strain genomes was not annotated in MGD. Encouragingly, we have thousands of additional gene function hypotheses for phenotypic experiments, suggesting rich grounds for novel gene function discovery. To aid with searching literature evidence for the most plausible candidate, we built a novel Literature-Based Discovery (LBD) framework. Finally, a web portal was developed to support easy browsing of all of our results.

## Materials and Methods

### Mouse strains

We used whole genome data of 48 inbred mouse strains in our analysis: C57BL/6J, 129P2/OlaHsd, 129S1/SvImJ, 129S5/SvEvBrd, AKR/J, A/J, B10.D2-H2<d>/n2SnJ, BPL/1J, BPN/3J, BTBR T<+> Itpr3<tf>/J, BUB/BnJ, BALB/cJ, C3H/HeJ, C57BL/10J, C57BL/6NJ, C57BR/cdJ, C57L/J, C58/J, CBA/J, CE/J, DBA/1J, DBA/2J, FVB/NJ, I/LnJ, KK/HlJ, LG/J, LP/J, MA/MyJ, NOD/ShiLtJ, NON/ShiLtJ, NOR/LtJ, NU/J, NZB/BlNJ, NZO/HlLtJ, NZW/LacJ, P/J, PL/J, RF/J, RHJ/LeJ, RIIIS/J, SEA/GnJ, SJL/J, SM/J, ST/bJ, SWR/J, TALLYHO/JngJ, RBF/DnJ, MRL/MpJ (Supplementary Table 1). Our reference strain is C57BL/6J (the “Black 6” mouse) and our reference genome assembly for all variant calls is GRCm38/mm10.

### Phenotype data

#### Inbred mouse phenotype data

Mouse Phenome Database (MPD)^8^ is a repository of experimental phenotypic data measured on various mouse inbred strains in different laboratories. We downloaded the animaldatapoints.csv.gz, strainmeans.csv.gz, and ontology_mappings.csv files on March 28, 2022 from https://phenome.jax.org/downloads. We kept all experiments *e* when at least 6 of our 48 sequenced strains were phenotyped, and the reported trait is relevant for genetic dissection (e.g., we removed experiment MPD:36680 reporting “age of mice at testing”).

In addition, 14 out of our 48 sequenced strains had at least 1 strain-specific, naturally occurring, spontaneous phenotype annotated in MGD^4^ (e.g., A/J mice spontaneously develop cochlear hair cell degeneration). For each phenotype, we labeled strains annotated with the phenotype 1 and strains not annotated with the phenotype 0. These phenotypes were added to experiments (Figure 1).

#### Phenotype separation boundaries

For each experiment *e*, we sought a phenotype separation boundary *b* that separates strains into two groups based on their phenotypic values. All strains used in the study are considered for the purpose of binarization, including strains that we do not have genomes for. Most MPD experiments reported biological replicate values, but some reported only the average and the standard deviation across replicates. We handle these two cases separately:

##### Experiments with replicate values

To remove any outliers, we discarded the top and bottom 25 percentile biological replicates for strains with more than 5 replicates. The strains were sorted by the average of the remaining biological replicates. Any boundary grouping that sorted strains into “left” vs “right” was recorded as separable if the following conditions were met: (1) the maximum replicate value in the “left” group is smaller than the minimum replicate value in the “right” group (2) the difference between the two values is greater than twice the average standard deviation of the two strains^11^ (Supplementary Figure 1).

##### Experiments without replicate values

The strains were sorted by the reported average of the biological replicates. For every strain in the sorted order, we recorded a phenotype separation boundary between the strain and the next adjacent strain if: (1) the strain’s average value plus its standard deviation was less than the next strain’s average value minus its standard deviation; (2) the difference between these two values is greater than twice the average standard deviation of the two strains.

This resulted in each experiment having zero, one, or multiple possible phenotype separation boundaries. Each boundary was considered independently in candidate gene matching.

### Genotype data

#### Whole genome variant calling

The raw reads of 48 strains were trimmed, filtered, and aligned to the reference genome (C57BL/6J, GRCm38/mm10) using Burrows-Wheeler Aligner (BWA)^12^ with default settings. Then the reads were realigned and recalibrated around indels using GATK(v3.6) ^13^ with default parameters. Single nucleotide polymorphisms (SNPs) and insertion or deletions (indels) discovery were performed using BCFtools (v1.9)^14^ command: “bcftools mpileup -a DP,AD,ADF,ADR,SP,INFO/AD -E -F0.25 -Q0 -p -m3 -d500”. Joint calling was performed on 47 strains (against the reference C57BL/6J strain) using: “bcftools call -mv -f GQ,GP”. Indels were then left-aligned and normalized using: “bcftools norm -d none -s -m+indels”. Low quality variants were filtered out using: “bcftools view -i ‘MIN(FMT/DP)>3 & MIN(FMT/GQ)>20’” and “bcftools filter -g3 -G10”.

#### Strain-specific variant filtering

We used Phred quality scores^15^ to measure variant call quality per strain. We required that the most confident call be at least 100 times more likely than the second most confident call (i.e., Phred score difference of at least 20). We marked the variant call for the strain as inconclusive if the gap was smaller than 20. While inbred mice are bred to homogenize as many of their alleles as possible, we kept any high-quality heterozygous calls as heterozygous.

#### Comparing and supplementing our sequencing data with a published dataset

Mouse Genome Project (MGP)^16^ publishes various mouse genetic variations including SNPs and Indels from 37 of our 48 inbred strains. Using their FTP site, we downloaded their SNP and Indels VCF on March 04, 2021 (ftp://ftp-mouse.sanger.ac.uk/REL-2004-v7-SNPs_Indels/mgp_REL2005_snps_indels.vcf.gz). All variants included had already gone through quality filtering. We applied the same Phred quality score filtering to find only the conclusive calls for each strain. After comparing the two sets, we replaced our inconclusive calls with MGP’s conclusive calls.

#### Annotating variants with Ensembl VEP

We annotated each variant using Ensembl Variant Effect Predictor (VEP)^17^. VEP tags each input variant with expected consequences of all relevant transcripts. We downloaded the newest version 102.0 for mm10 on February 24, 2021. To remove any ambiguous calls due to alignment orientation, we ran VEP using both left and right alignments on our mouse strain variant call format (VCF) files. We performed left alignment using GATK’s LeftAlignAndTrimVariants tool and the mm10 Fasta sequence file downloaded on March 01, 2021 from: https://hgdownload.soe.ucsc.edu/goldenPath/mm10/bigZips/chromFa.tar.gz. GATK’s left alignment tool requires index and dictionary files for these Fasta files. We ran samtools faidx command to make the index files. We downloaded picard.jar version 2.25.4 from https://broadinstitute.github.io/picard/ on March 06, 2021 and ran the following command to make the dictionary files:

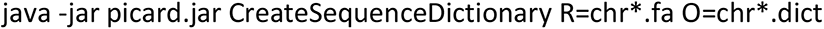

Then, using the sequence Fasta files with their index and dictionary files and our VCF files as inputs, we ran the following command to make left aligned VCF files:

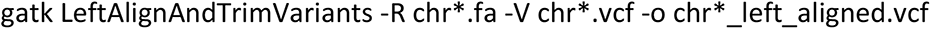

We used the VEP Docker image to annotate our VCFs. We ran VEP once with our GATK left aligned VCF for left aligned annotations and once with our original VCF with VEP’s – shift_genomic flag set to True for right aligned annotations.

#### Gene set

We built a gene set from GENCODE VM32 known genes downloaded from the UCSC Table Browser^18^ on March 01, 2021. We removed all transcripts that do not have a coding sequence that starts with a start codon (ATG) and ends with a stop codon (TAA, TAG, and TGA).

#### Genes with high-impact variants

We kept a variant if and only if it was tagged with a high-impact consequence to any transcript in our gene set by VEP using both left and right alignments (Supplementary Figure 2). High-impact consequences defined in the VEP documentation are: transcript ablation, splice acceptor variant, splice donor variant, stop gained, frameshift variant, stop lost, start lost, and transcript amplification. VEP annotates each variant independently. To account for adjacent SNPs affecting the same codon, we performed a post processing step to check if the final amino acid resulting from the effect of all SNPs would indeed result in a high-impact change. Finally, we removed any high-impact variants in predicted genes, RIKEN genes, Olfactory and Vomeronasal genes. We represented all genes using NCBI gene IDs.

### Building candidate sets

For each separable boundary *b* derived from the phenotypic experiment *e*, we searched for concordant genotype separations for each highly-impacted genes. We merged the genotypes per strain per gene *g* across transcripts and variants (Supplementary Figure 3):

*homozygous reference strain*: has no homozygous or heterozygous high-impact alternate allele.

*homozygous alternate strain*: has at least 1 homozygous high-impact alternate allele.

*heterozygous alternate strain*: has at least 1 heterozygous high-impact alternate allele and no homozygous high-impact alternate allele.

*inconclusive strain*: has at least 1 inconclusive gap or low-quality allele that may be high-impact and has no homozygous or heterozygous high-impact alternate allele.

*unsequenced strain:* not one of our 48 sequenced strains.

A candidate gene *g* is a match for experiment *e* with boundary *b*, if the following conditions are met: (1) all homozygous alternate strains and all homozygous reference strains are separated by *b*, (2) both sides of *b* have at least one homozygous strain, (3) there are at least 6 total homozygous strains between the two groups, and (4) all heterozygous alternate strains are on one side of *b*. Any unsequenced strains or inconclusive strains can appear on either side of *b* (Supplementary Figure 3). All matching experiment *e*, boundary *b*, gene *g*, (*e, b, g*) triplets form our trait-specific candidate set.

### Validating against known causative gene-phenotype relationships

We searched for known gene-phenotype relationships in our candidate set as proof of principle to our approach. We used Mammalian Phenotype (MP)^5^, an ontology of phenotypes observed in mice, to find matches.

For each experiment *e*, we obtained one or more MP terms Φ_*e*_, denoting the phenotype measured in *e* from the experiment source (MPD or MGD). For each gene *g*, we obtained a set of zero or more MP terms Φ_*g*_, denoting the phenotype annotations supported by single gene knockouts from MGD. Φ_*g*_ for all available genes were downloaded on April 26, 2021 through http://www.informatics.jax.org/downloads/reports/MGI_PhenoGenoMP.rpt.

For each trait-specific candidate (*e, b, g*) we generated, we first asked whether |Φ_*e*_ ∩ Φ_*g*_| > 0, which implies a known causal relation between *e* and *g*. All such matches are listed in Table 1, and they represent cases where our candidate gene has already been shown to underlie the measured trait in experiment *e*.

**Table 1.**
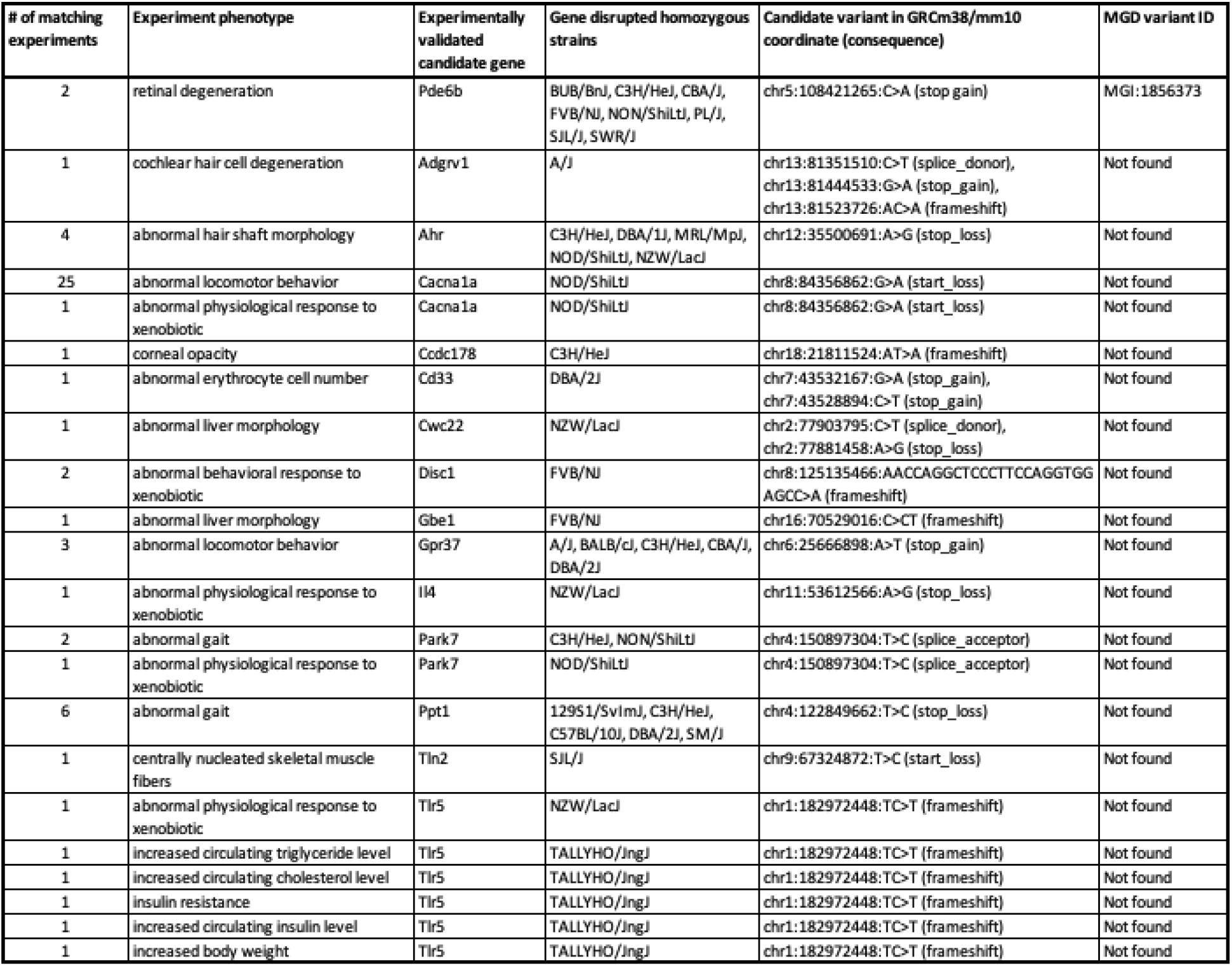
Known gene functions successfully rediscovered by our approach. In all shown cases, the candidate gene listed is already known to cause the experimental phenotype. However, only in the first (*Pde6b*) is the high-impact variant we found documented in MGD. See all details at aimhigh.stanford.edu (Supplementary Figure 6).

MP terms are organized in a hierarchical structure where a lower, more specific term (i.e., descendant terms, *D*(Φ)) is implied by its higher, more general term. Since the experiments measure a spectrum of phenotypes across different strains, they are often tagged with a more general term than the gene. As an example, the experiment MPD:22970 is tagged with “abnormal mean corpuscular volume (MP:0000226)”, whereas its candidate gene *Foxp3* is tagged with the more specific descendant term “increased mean corpuscular volume (MP:0002590)”. To benefit from this hierarchical structure, we also listed additional positive controls |Φ_*e*_ ∩ *D*(Φ_*g*_)| > 0 in Table 2.

**Table 2.**
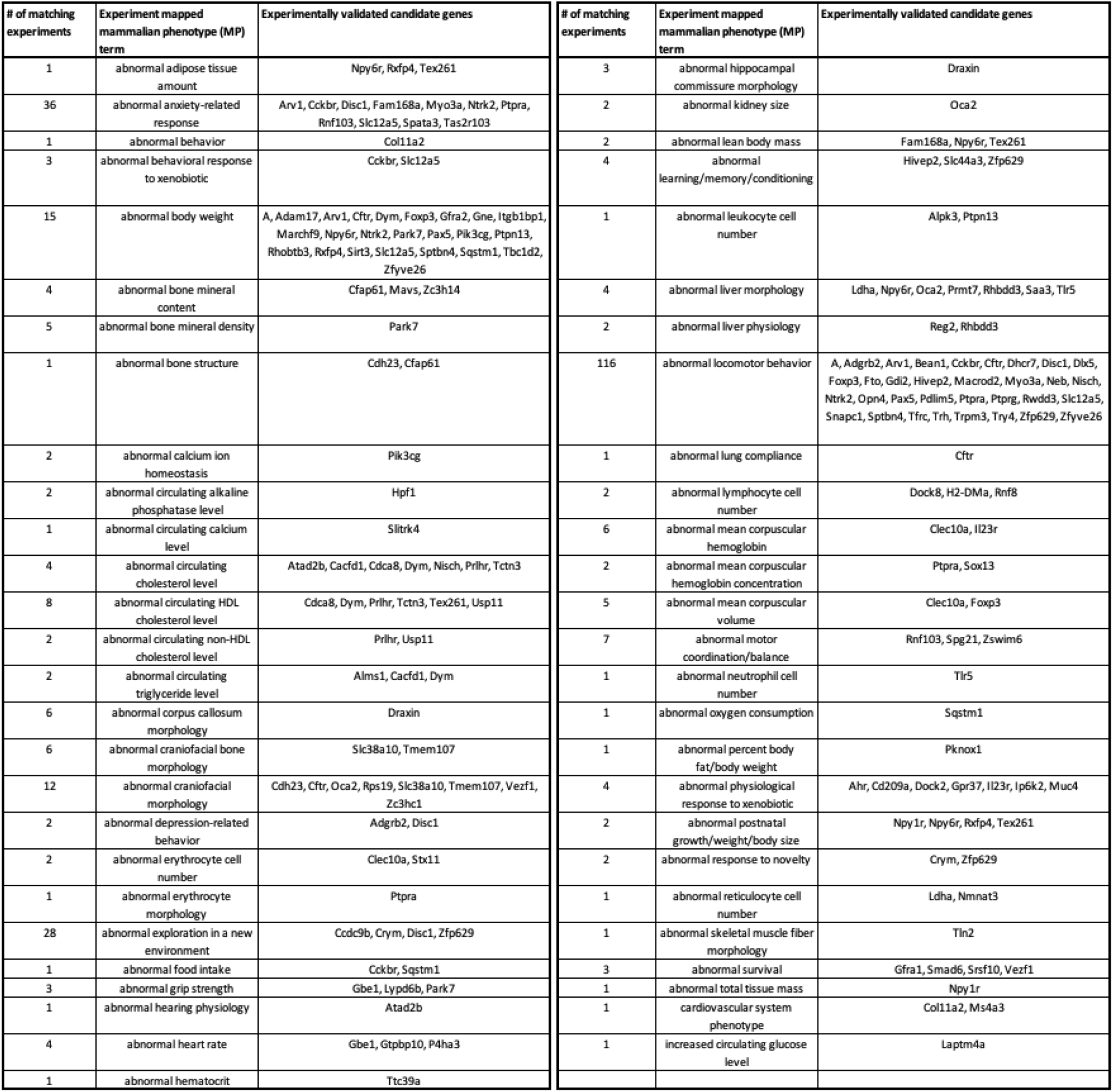
Additional known gene functions successfully rediscovered by our approach. In Table 1, the gene annotated function was identical to the experimental term. Here, the gene annotated function is a descendant term of the experimental term. For example, the experiment was tagged with abnormal body weight while its candidate gene, *Pax5*, is annotated with decreased body weight. Because of the large number of matches, we aggregated results by the experimental term. See all details of each match at aimhigh.stanford.edu (Supplementary Figure 6).

### Literature-based discovery (LBD) classifier

#### Motivation

Encouragingly, over 200 gene-phenotype relationships from our trait-specific candidate sets were previously known (Figure 1). This indicates our method is effective in discovering true relationships. At the same time, thousands of our gene-phenotype hypotheses currently have no reported causative relationship, suggesting many exciting hypotheses yet to be discovered.

Some of our experiments yielded very few novel candidate genes. Table 3 holds a sample of experiments with fewer than 10 candidate genes.

**Table 3.**
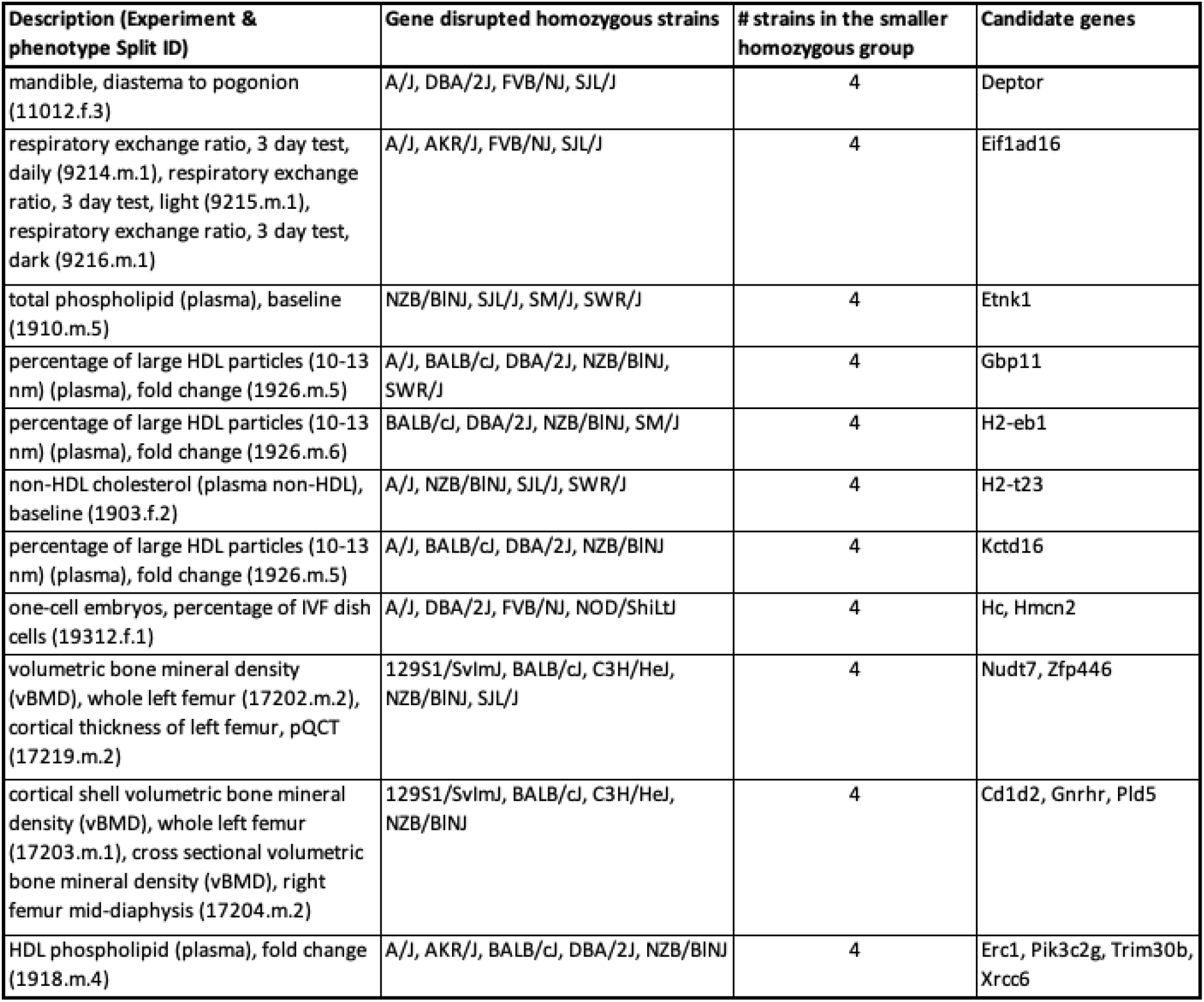
A sampler of novel hypotheses with few candidate genes. Many of our novel hypotheses have very few matching candidate genes per experiment. A handful is shown here. All matches with fewer than 10 candidate genes per experiment are available at aimhigh.stanford.edu (Supplementary Figure 6).

The remaining experiments each have as many as 103 candidate genes. To build a small set with compelling evidence for further investigation, a researcher would need to carefully read literature on all candidate genes we matched in each experimental context. To mimic the process, we devised a supervised Machine Learning (ML) approach to predict which gene-phenotype relationships would become true in a few years based on existing literature evidence. Briefly, we trained a ML classifier to learn patterns from literature up to year *T*_*0*_ to be predictive of gene-phenotype relationships that were discovered later at some time *T*_*1*_. We used the trained classifier to prioritize the most promising novel candidates for each mouse inbred strain experiment *e* (Supplementary Figure 4A). Exciting examples are shown in Table 4.

**Table 4.**
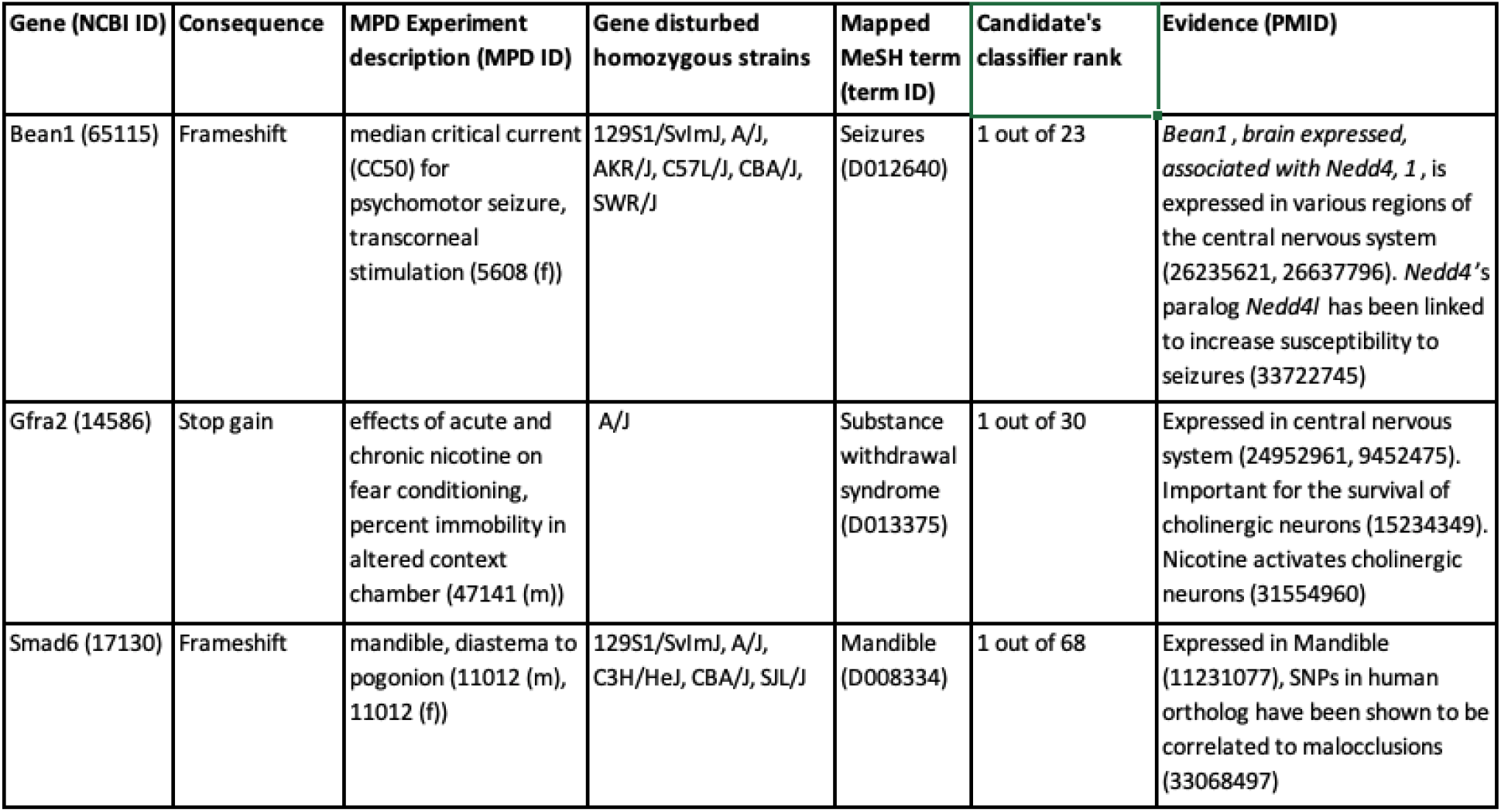
A sampler cases where our classifier (Figure 1B) singles out biologically exciting candidate genes. All novel hypotheses, including our classifier’s top-ranking genes and supporting literature are shown at aimhigh.stanford.edu (Supplementary Figure 6).

#### MeSH-publication relationships

We used PubMed, a database of over 33 million peer-reviewed biomedical publications, as our literature source. They key topics mentioned in PubMed papers are manually indexed using the Medical Subject Heading (MeSH) ontology^19,20^. These topics include functional annotations but not gene mentions. Similar to MP, MeSH is also organized in a hierarchical structure with descendant terms are implied in the ancestor terms. For example, “Microcephaly (D008831)” is a descendant of “Disease (D004194)”. MeSH term annotations per paper (in PubMed ID, or PMID) were downloaded from https://ftp.ncbi.nlm.nih.gov/pubmed/baseline/ on April 12, 2022.

#### Gene-publication relationships

We downloaded mentions of genes in each PubMed abstract from Pubtator Central^21^. Pubtator Central^21^ is a natural language processing (NLP) tool that tags PubMed papers with recognized entities such as genes. The mappings between genes (in NCBI ID) and abstracts (in PMID) were downloaded using https://www.ncbi.nlm.nih.gov/research/pubtator-api/ on April 12, 2022.

#### Publication graph

To encode relatedness, we built a publication graph where a node is either a MeSH term or a gene. We extended an undirected edge between any two nodes (MeSH-gene, gene-gene, or MeSH-MeSH) if there is at least one paper that mentions them together (Supplementary Figure 4B).

#### Setting up the learning task

Our supervised learning task requires a set of labeled examples (positive or negative) and a learning algorithm. Each example is represented by a set of scalar values (i.e., features).

#### Single-gene knockout positive set

We trained our ML classifier on positive examples derived from papers that describe the phenotypic impact discovered in single-gene mouse knockout (KO) experiments. A single-gene mouse knockout paper is tagged as a knockout of exactly one gene in MGD and tagged with a MeSH term, “Mice, Knockout (D018345)” in PubMed.

We downloaded these papers from MGD’s MouseMine API (https://www.mousemine.org/mousemine/begin.do) on April 12, 2022. Each single-gene *g* KO paper was tagged with one or more phenotype MeSH terms *m* to build a set of (*g, m*) pairs. Given all the literature evidence at time *T*_*0*_, we aim to predict which (*g, m*) pairs would be discovered at some later time *T*_*1*_ (Supplementary Figure 4A, C).

#### Building retrospective single KO train and test sets

To propose plausible (*g, m*) pairs for future discovery, we used the open discovery “ABC” method^22,23^. For a node A in our graph, we found nodes, C, that are linked at *T*_*0*_ by one or more intermediary node B (i.e., A-B and C-B edges) to A but have no direct A-C link. All A-C links are candidates for discovery at *T*_*1*_. To perform retrospective analysis, we built a time-specific publication graph using papers published up to *T*_*0*_ and compared it against a publication graph at *T*_*0*_ *+ 5* (Supplementary Figure 4C).

We used two publication graphs to train our model: the 2010 graph (i.e., *T*_*0*_) and the 2015 (i.e., *T*_*0*_ *+ 5*) graph. We found (*g, m*) pairs mapped to single-gene knockout publications published between (2010, 2015]. We discarded a *(g, m*) pair if any of the following was true: (1) *g* and *m* were directly linked in the 2010 graph, (2) either *g* or *m* were not found in the 2010 graph, or (3) *g* and *m* did not share an intermediate node in the 2010 graph. The remaining pairs are our positive examples.

We added negative examples (i.e., (*g, m*) pair not discovered by 2015) for each MeSH term *m* that is part of at least 1 positive example. A (*g, m*) pair is a negative example if the following conditions were met: (1) *g* is a mouse protein-coding gene from our gene set found in the 2010 graph, (2) *g* shares at least 1 intermediate node with *m* in the 2010 graph, and (3) *g* does not have a direct link to *m* in the 2010 and the 2015 graph.

We repeated the same operations with the 2015 graph and the 2020 graph to test our algorithm (Supplementary Figure 4A, Figure 2A).

**Figure 2.**
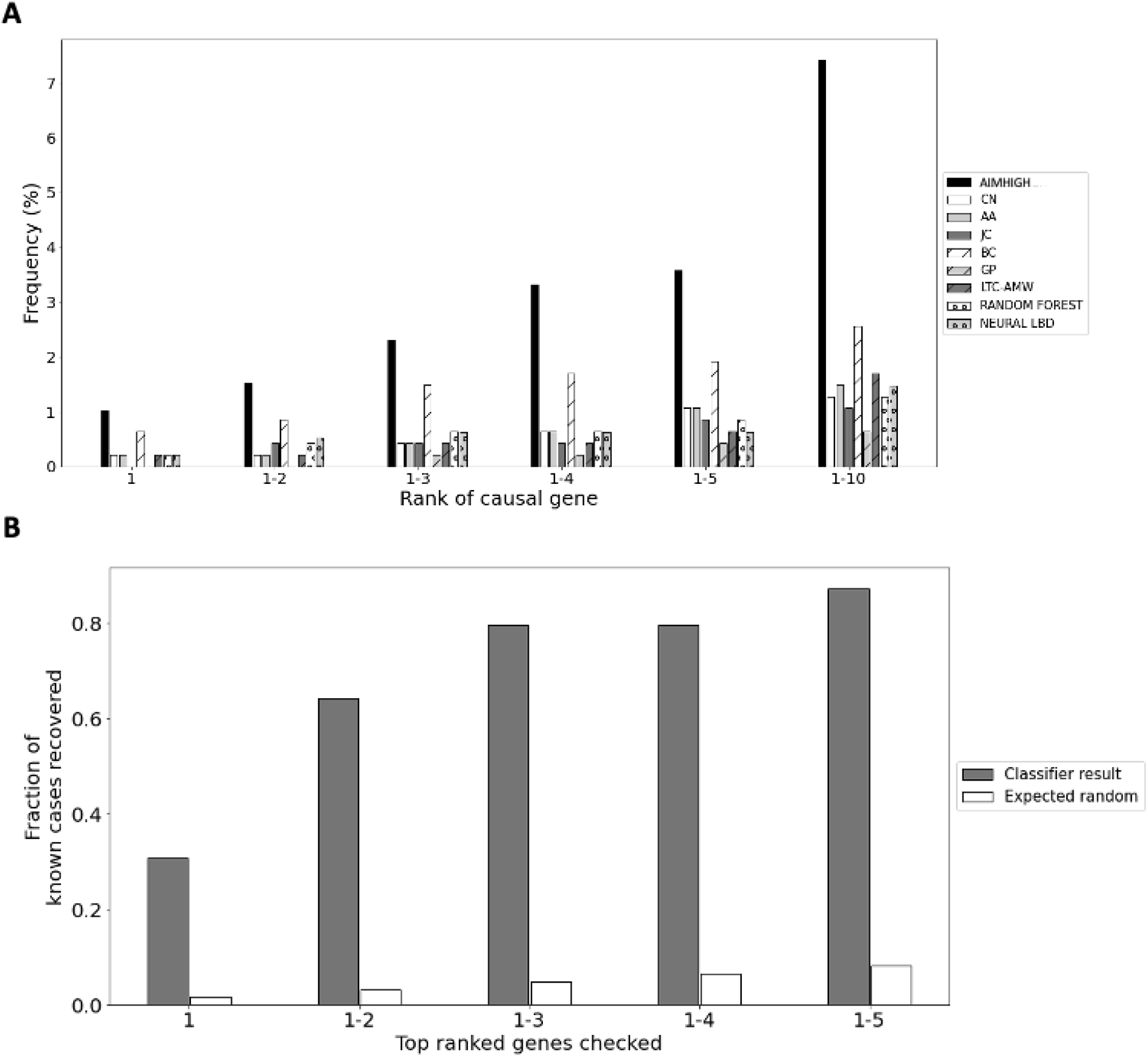
Predicting which gene function hypotheses would be proven over a 5-year period. **(A)** Our machine learning classifier performance on an open discovery set built from mouse single gene knockout papers, where on average out of 5515.8 candidate genes, only 2.4 genes are verified over the next 5 years (see Methods). While this task is difficult for all 8 approaches, AIMHIGH clearly improves the state of the art. **(B)** By matching phenotypic separation with genotypic separation (Figure 1 and Supplementary Figure 3), we greatly reduce the search space for each trait. When we apply our classifier to known cases in our set (Tables 1 and 2) with at least 10 candidate genes discovered over the same 5 year period, we see that the causative gene ranks in top 3 in nearly 80% of cases (compared to 4.8% if 3 candidate genes are picked at random), suggesting that reading (or even directly testing) a handful of our top-ranked novel candidate genes may be highly effective in discovering novel biology. See Methods for more on all 8 tested approaches.

Open discovery resulted in thousands of negative examples for each positive example (see results), making ranking the positive examples on top a challenging task.

#### Building rediscovered test set and candidate hypothesis prediction set

Because we matched genomic signatures to phenotypic separations, our actual sets of candidate genes are orders of magnitude smaller. To assess the performance of our algorithm on our set of novel hypotheses, we applied it to our set of rediscovered hypotheses (Supplementary Figure 4A, Figure 2B). Then we applied the method to our novel candidates to prioritize the most exciting hypotheses (Table 4).

Since PubMed uses MeSH terms to annotate papers, we mapped each phenotypic experiment *e* to zero or more MeSH terms *m* that denote the phenotype measured in *e*.

From our rediscovered test set (Tables 1 and 2, see Figure 1), we selected a set of experiment *e* and boundary *b* that has at least 10 candidate genes. From this set, we searched for positive (*g, m*) pairs where *g* and *m* only shared intermediate nodes in 2015 but were directly linked in the 2020 graph. All non-positive (*g, m*) pairs from the same experiment *e* and boundary *b* were labeled negative (Supplementary Figure 4A, Rediscovered single KO test set).

Finally, our candidate hypothesis prediction set contained novel (*g, m*) pairs that as of July 2022 have no known relationship in MP or MeSH (Supplementary Figure 4A, Candidate hypothesis prediction set).

#### Classifier features

We used the publication graph to provide the classifier a set of features (i.e., clues) that represent each gene *g*, MeSH term *m*, (*g, m*) pairs in our sets.

We used the appropriate time-specific publication graphs to build these features. For example, we used the 2010 graph to build features to predict relationship status in 2015.

We first define these terms for node *u and v* (MeSH term or gene) in our publication graph to help explain each feature:

Neighbors of u: *N*(*u*) = *set of* nodes directly connected to u

Degree of u: *d*(*u*) = |*N*(*u*)|

Publications of u: *p*(*u*) = publications that mention *u*

Weight of an edge connecting *u* and *v*: 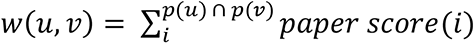

Paper score of *i*: 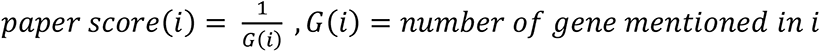

The following features are used in our classifier:

##### Degree of the gene

*d*(*g*) (number of nodes directly connected to *g*)

##### Common neighbor^24^

|*N*(*g*) ∩ *N*(*m*)| (number of nodes directly connected to *g* and *m*)

##### Adamic Adar^25^

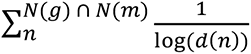 (sum over the inverse log of number of nodes directly connected to shared neighbors of *g* and *m*)

##### Preferential attachment^24^

*d*(*g*) *\* d*(*m*) (number nodes directly connected to *g* times number nodes directly connected to *m*)

##### Gene publications

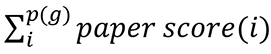

##### Human ortholog link

The weight of the edge between m and *g*’s human ortholog. Orthologous relationships are defined by Ensembl 102, latest version with mm10, downloaded from http://nov2020.archive.ensembl.org/biomart/martview/ on September 3, 2021. If there were multiple orthologs then the average value is used, and if there is no direct edge or there is no human ortholog in the graph then we assign (-1).

##### BITOLA confidence^26^

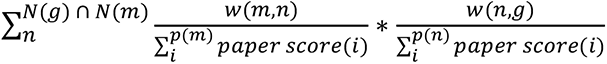

##### Reverse BITOLA confidence

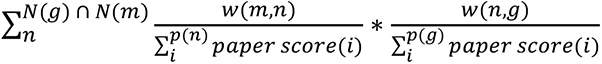

##### MeSH term’s gene neighbors

the number of genes directly mentioned with *m*.

##### Mean expression

find top 5 anatomical entities (i.e., descendant MeSH terms of “Anatomy (D000715)” such as “Brain (D001921)”) that are most often co-mentioned with *m*. Then take the average of the Adamic Adar scores between the top 5 anatomical entities and *g*.

#### Classifier model

We used a Gradient Boosting Tree Classifier^27^ to assign scores to each example. The score is a number ranging from 0 to 1, and a higher number indicates the example is more likely to be positive. Gradient Boosting Trees is a supervised method which means it is presented with a large number of labeled examples (i.e., training set) and it learns to distinguish the two labels by minimizing the final entropy. It is an ensemble method that involves a series of decision trees that are iteratively constructed to optimize the performance on hard-to-classify training examples. In each iteration, any examples that were misclassified (a positive example classified as negative, or vice versa) have a higher chance of being selected in the next iteration so that the decision trees can learn to classify them correctly. We used a Gradient Boosting Tree Classifier implementation contained in the python package sci-kit learn v0.18.1.

The Gradient Boosting Tree function F of input x in stage m (value of the output at the stage m) is calculated using the formula below:

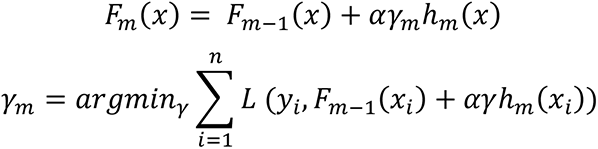

where γ_m_ is a value that is chosen to minimize the loss function and gets multiplied to a decision tree h_m_. α is a hyperparameter that represents the learning rate which controls how much each tree contributes to the output.

#### Existing methods and baselines

We compared our classifier’s performance against 8 existing methods. We used 6 non-learning-based methods: Common neighbor (CN)^24^, Adamic Adar (AA)^25^, Jaccard coefficient (JC)^28^, BITOLA confidence (BC)^26^, Gene publications (GP), and Linking Term Count with Average Minimum Weight (LTC-AMW)^29^.

We also used 2 learning-based methods: Random Forest approach described in Kastrin et al.^24^ and neural LBD by Crichton et al.^22^. The non-learning-based methods’ scores were calculated for the retrospective single KO set, and the learning-based methods were trained and tested using the same inputs as our LBD classifier.

#### Model evaluation

Performance is measured by how often a gene with a causative relationship to the measured trait is ranked at or near the top.

We have a set of candidate genes per each phenotype MeSH term *m* in the retrospective single KO test set and per each experiment *e*, MeSH term *m*, and boundary *b* in the rediscovered test set. We reported the rank of the causative gene *g* within the group of candidate (positive and negative) genes (Figure 2).

#### Explaining our predictions using supporting literature evidence

To facilitate follow up researcher evaluations of our highest-ranking hypotheses, we sought to highlight each pair of phenotype-candidate gene with a curated list of papers that helps support our predictions.

We first selected up to three most relevant intermediate nodes (MeSH term or gene) linking the phenotype MeSH term and the hypothesis gene using the BITOLA confidence score. Then, we reported the most recent paper that mentions the evidence terms and the hypothesis gene. If there are multiple papers published in the same year, we picked the paper with the highest paper score (Supplementary Figure 5).

## Results

The workflow of our results is shown in Figure 1.

### Phenotype data

#### Inbred mouse phenotype data

11,783 measurements were downloaded from MPD. We kept 4,939 measurements after removing any experiment that did not use at least 6 of our sequenced strains (Supplementary Table 1). 610 of them were not expected to be genotypically relevant (e.g., reporting mouse age at time of experiment). The remaining 4,329 measurements were used for candidate gene selection. Along with 49 additional experiments we created for presence/absence trait MGD reported across 14 of our strains (see Methods).

#### Phenotype separation boundaries

4,110 suitable MPD cases were reported in biological replicate values and the remaining 219 cases were only reported in average, standard deviation, and number of replicates used.

729 suitable MPD cases had at least one phenotypically separable boundary (Supplementary Figure 1). 49 experiments we added were separated by the absence/presence of the MP term. In total, we found 1,133 phenotype separation boundaries in 778 experiments.

### Genotype data

#### Whole genome variant calling

We kept 23,230,042 SNPs and indels among 48 mouse inbred strains after discarding any variant that failed to meet the quality threshold (Supplementary Figure 2).

#### Strain-specific variant filtering

Using Phred-quality scores to filter strain-specific calls, 1.51-12.1% of SNPs and indels had inconclusive calls per each of the 47 strains (excluding the reference strain). Through generations of sibling mating, inbred mice are expected to have largely homogenized alleles. Indeed, we found that only 0.18-0.75% of our variant calls were high quality heterozygous.

#### Comparing and supplementing our sequencing data with a published dataset

We found that 742,480,455 or 99.9% of our 743,170,690 conclusive allele calls in the overlapping 37 strains were supported by the MGP data (732,135,280 or 98.5% identical and 10,345,175 or 1.4% inconclusive by MGP).

5,812,957 (23.3%) of 24,925,741 calls we called inconclusive were called conclusively by MGP. After supplementing with MGP, our final calls in 47 strains include 817,092,836 (74.8%) homozygous reference, 232,498,308 (21.3%) homozygous alternate, 6,039,276 (0.6%) heterozygous, and 36,181,554 (3.3%) inconclusive calls.

#### Gene set

Our gene set contained 50,527 transcripts of 21,713 unique protein coding genes. In our downstream analysis, we only considered VEP’s variant consequences that affect the transcripts in our gene set.

#### Genes with high-impact variants

We identified 2,178 transcripts for 1,409 genes with at least 1 high-impact variant in at least one non-reference inbred mouse strain (Supplementary Figure 2). A high-impact variant can be either homozygous or heterozygous for one, or even two different high-impact alleles (e.g., 1/2 where 1 and 2 both represent high-impact alternate allele).

### Building candidate sets

Each of 1,133 separable phenotypic boundaries, was compared to our 1,409 highly-impacted genes to build a candidate gene-phenotype set (Supplementary Figure 3 and Methods). A total of 906 phenotype separations in 718 unique experiments had at least 1 candidate gene with an average of 17.53, a median of 12, a minimum of 1, and a maximum of 103 candidate genes per phenotype separation.

### Previously reported causative gene-phenotype relationships

We downloaded 296,254 gene markers to phenotype relationships (33,010 markers and 10,741 MP terms) from MGD^4^. We mapped the gene markers to NCBI gene IDs in our gene set and kept only gene-phenotype relationships unambiguously mapped to 1 gene. The resulting set included 190,623 single gene-phenotype relationships among 10,054 MP terms and 12,551 protein-coding mouse genes.

### Candidate genes previously linked to the measured phenotypes

#### Exact Match

We found 22 gene-phenotype relationships where the gene and the experiment were tagged with the same MP term suggesting our approach (Figure 1) picked the correct gene to explain the experiment (Table 1). Interestingly, in only one of the cases, the high-impact variant we found had an MGD record of causing the measured phenotype. In all other cases, depending on the completeness of MGD, we may be the first to report the strain(s) specific potentially causative variant(s).

For example, DBA/2J mice had elevated red blood cell counts compared to 10 other strains (MPD:22903). 33 candidate genes fit this phenotype separation and only 1 gene, *Cd33*, is known to cause the same phenotype, Abnormal erythrocyte cell number (MP:0001586). DBA/2J uniquely has a stop gain mutation (GRCm38/mm10 chr7:43528894) in *Cd33* that may explain abnormally high red blood cells in this strain but missing in MGD.

Full details of all cases are available at aimhigh.stanford.edu (Supplementary Figure 6).

#### Descendant match

There were 190 gene-phenotype relationships where the gene was mapped in MGD to a more specific (descendant) MP term of the experiment phenotype. These are shown in Table 2. Only one high-impact variant is known in MGD, and other variant-phenotype relationships may be novel. Full details of all cases are available at aimhigh.stanford.edu (Supplementary Figure 6).

As an example, MPD:1903 measured non-HDL cholesterol levels in 14 mouse strains and found abnormally high non-HDL cholesterol level in NZB/BINJ mice. This experiment is tagged with Abnormal circulating non-HDL cholesterol level (MP:0020151). Among 41 candidate genes that contained a unique high-impact variant in NZB/BINJ, *Prlhr* is the only gene annotated with a matching descendant phenotype term, Increased circulating LDL cholesterol level (MP:0000182). There is a potentially causative stop gain mutation (GRCm38/mm10 chr19:60467194) in NZB/BINJ not found in any other strains, and this variant is not recorded in MGD.

### Literature-based discovery (LBD) classifier

#### MeSH-publication relationships

We downloaded a vocabulary of 29,405 MeSH terms tagging 28,726,608 of 33,425,447 papers published between 1902 and 2022 for a total of 303,924,223 MeSH term *m*, publication *pub* (in PMID), (*m, pub*) pairs.

#### Gene-publication relationships

We downloaded 177,518 genes from 3,478 species mentioned in 6,661,514 papers published up to 2022, for a total of 14,632,626 gene *g*, publication *pub*, (*g, pub*) pairs.

#### Publication graph

The full 2022 graph used to featurize our candidate hypothesis prediction set (Supplementary Figure 4A) has 79,306,096 edges. Nodes have 48 median direct neighbors and the density of the graph, a ratio of number of edges present over the maximum number of possible edges in our graph, is 0.004.

#### Single-gene knockout positive set

We downloaded 117,438 papers that contain the “Null/Knockout” attribute and are linked to exactly 1 gene. 61,403 papers describe a phenotype measured indicated by tagged disease MeSH terms about 8,268 unique mouse protein-coding genes.

#### Building retrospective single KO train and test sets

For the retrospective single KO train set, the labels of each gene *g*, MeSH term *m*, (*g, m*) pairs (i.e., positive for relationships defined by a single-gene knockout paper and negative for unknown relationships) were derived from single-gene mouse knockout papers that were published between 2011 and 2015, inclusive (Supplementary Figure 4A). This set consisted of 1,619 positive edges and 2,516,571 negative edges for 568 MeSH phenotypes (2.9 positive and 4,421.7 negative mouse genes per MeSH term on average).

For the retrospective single KO test set, we used single-gene mouse knockout papers that were published between 2016 and 2020, inclusive. This set consisted of 1,130 positive edges and 2,586,932 negative edges for 469 MeSH phenotypes (2.4 positive and 5515.8 negative mouse genes per MeSH term on average).

#### Building rediscovered test set and candidate hypothesis prediction set

We mapped 906 phenotype groupings to 170 unique MeSH terms. 47 phenotypes such as “KRTAP14, spectral counts, hair proteomics (MPD: 49233)” were not mapped because they lacked a suitable MeSH term.

In our rediscovered test set, we found 41 positive (*g, m*) pairs with at least 10 candidate genes whose direct relationship was established after 2015, with a total of 898 negative pairs. Per experimental boundary, there were 24.1 candidates and only 1.1 of them came true after 2015 on average. Note that our phenotype-genotype matching (Figure 1 and Supplementary Figure 3) results in a significantly reduced candidate hypotheses compared to the literature-based, open discovery “ABC” method^22,23^ used in the above sets.

In our candidate hypothesis prediction set, we found 13,652 novel (*g, m*) pairs whose experiment had at least 10 candidate genes and no known relationship to the measured trait. Per experimental boundary, there were an average of 24.7, a minimum of 10 and a maximum of 103 candidate genes.

#### Classifier performance

Open discovery is useful in automatically generating hypotheses^10,22^, but this approach resulted a very large candidate hypotheses set (thousands of negative examples for each positive example). In Figure 2A, we show that on this exceptionally difficult task, our AIMHIGH classifier beats 8 existing methods. For example, in the retrospective single KO test set it ranks the positive edge in the top 10 among over 5,000 candidate genes in 34 out of 469 MeSH terms (7.4%) compared to 2.6% by the next best method, BITOLA confidence^26^.

We tested AIMHIGH on a secondary test set to assess how it would perform on a set with more realistic candidate sizes. Figure 2B shows this using our rediscovered test set. After 2015, predicted from the 2015 graph it ranked the positive gene among top 3 in 79.5% of the cases compared to the expected 4.8% if we ranked all hypotheses randomly (Figure 2B).

### Novel hypotheses

#### Novel cases with few candidates

1,092 gene-phenotype relationships from our inbred strain analysis did not have a known explanation and had fewer than 10 candidate genes per phenotype boundary. In such cases, all candidates can be reviewed closely in search of enticing hypotheses for testing. 17 such gene-phenotype relationships with at least 4 homozygous allele strains in the smaller phenotype group are shown in Table 3. Full details for all 1,092 relationships are available on at aimhigh.stanford.edu (Supplementary Figure 6).

For example, 7 strains showed low respiratory exchange ratio (RER) compared to 6 other measured strains (MPD:9214, MPD:9215, MPD:9216). RER is the ratio between carbon dioxide produced through metabolic process and oxygen consumed. Lower RER indicates higher fitness level and higher muscle’s ability to get energy^30^. Only 1 candidate gene was found to match this phenotype split, *Eif1ad16, eukaryotic translation initiation factor 1A domain containing 16*. Among the 7 strains in the lower group, 4 strains (A/J, AKR/J, FVB/NJ, and SJL/J) had a homozygous frameshift mutation, 2 strains (BTBR T<+> Itpr3<tf>/J and C3H/HeJ) had inconclusive calls, and 1 strain (129S1/SvImJ) had a heterozygous mutation in this gene. All strains in the higher group had homozygous reference alleles except LP/J which had an inconclusive call. *Eif1ad16* is currently not associated with any phenotypes in MGD. However, the eukaryotic translation initiation factor family are key regulators of translation initiation, and genes in this family have been suggested to initiate protein synthesis during recovery after resistance training^31^. We hypothesize that the strains high-impact variant in *Eif1ad16* have faster muscle protein metabolism indicated by lower RER.

### LBD classifier’s top-ranked predictions

Inspired by our rediscovered test set, we focus on the top 3 candidates predicted by AIMHIGH in all cases where 10 or more candidate genes exist. We found 750 most promising novel gene-phenotype hypotheses. A few examples are shown in Table 4 and discussed below.

MPD:5608 measured median current to trigger psychomotor seizure. C57BL/6J showed the highest resistance compared to 8 other phenotyped strains. 6 of the 8 strains were sequenced and all shared the same frameshift variant in *Brain expressed, associated with Nedd4, 1* (*Bean1*) gene. Our classifier ranked this gene above all other 22 other candidate genes. *Bean1* is expressed in the central nervous system^32,33^, and the paralog of *Nedd4* has been linked to susceptibility to seizure^34^.

A fear test after exposing mice to nicotine showed that A/J mice were less sensitive to nicotine effects compared to 7 other strains (MPD:47141). 30 candidate genes fit this phenotype split. Our classifier ranked *Gfra2* highest on the list. Promising evidence showed that cholinergic neurons are activated by nicotine^35^, and *Gfra2* is required for the survival of cholinergic neurons^36,37^.

MPD:11012 measured diastema to pogonion distance. C57BL/6J and C57BL/10J had longer distances than all other the measured strains. These two also were the only strains phenotyped that had the homozygous reference allele in *Smad6*. There were 68 candidate genes that fit this phenotype split, but the classifier ranked this gene highest. Encouragingly, MGD has recorded that *Smad6* is expressed in the mandible, and this gene has been shown to be correlated with malocclusion^38^.

Hundreds of similar leads can be found at aimhigh.stanford.edu (Supplementary Figure 6).

### Interactive web interface

aimhigh.stanford.edu holds an easy-to-use web interface for users to view all of our findings in detail. There are four webpages corresponding to Tables 1-4. See Supplementary Figure 6.

## Discussion

We present AIMHIGH, an automated approach (detailed in Figure 1) to suggest gene-phenotype testable hypotheses from community-contributed multi-strain phenotypic experiments and multiple strain genomes. By automatically finding one or more candidate gene whose high-impact variants split concordantly with the phenotypic measurement splits, we rediscovered hundreds of experimentally proven gene-phenotype relationships, validating our approach. Nearly all variants we highlighted in the context of these known gene functions are not found in MGD. More excitingly, we made thousands of potentially novel gene function hypotheses in experiments where we none of the matched candidates are already known to cause the measured phenotype.

We developed a machine learning approach that leverages existing literature to highlight a handful of most promising gene candidates among 10 or more such candidates. Our approach relies on relatedness of the millions of peer-reviewed papers indexed by PubMed and annotated by MeSH and PubTator Central. We started by improving the state of the art in literature-based open discovery, working with an extremely challenging set of thousands of candidate genes for every gene function hypothesis where only few were validated 5 years from our prediction time. We purposefully chose a relatively long time-period so that seemingly new discoveries will not simply be from those papers archived, or conference abstract announced at the time we make our predictions.

We then see again the power of our candidate building approach, which reduced the number of candidate genes per phenotype by two orders of magnitude compared to literature-based search. We show that even on the set without a known causative gene, one may only need to read on our top 3 highest ranked candidates per unexplained experiment to find the most plausible hypotheses.

Our goal throughout the screen is to avoid false positive predictions. Conservatively, we only screened for high-impact variants that are more likely to impact the protein function. Variants that can be left shifted or right shifted to provide an alternative, less impactful, interpretation are discarded, and more.

In the future, one can consider ways to extend our approach to lower impact variants such as non-synonymous substitutions (most of which have only a modest effect on gene function) and to structural variants (whose impact may often be high, but whose confident calling is challenging). Likewise, one can consider extending from the binary split we make into ones with more states, though the mapping of multi phenotypic groups to observed genomic changes becomes more challenging.

To encourage our colleagues to discover novel biology using our exciting predictions, we built a web portal at aimhigh.stanford.edu, which houses all of our rediscovered and novel predictions. Our code is also open sourced, so that anyone can rerun it as is, with new sequenced strains, additional phenotypic experiments, presence/absence traits or newly published literature strengthening our inference and deriving novel testable hypotheses.

## Data and code availability

Data and code will be available upon publication at https://github.com/bejerano-lab/AIMHIGH.git.

## Acknowledgements

We thank the members of the Bejerano laboratory, particularly Yosuke Tanigawa and Heidi Chen for technical advice and valuable feedback.

## Figures & Tables

**Supplementary Table 1.** 48 strain names used in this project along with their MGI strain ID, Jackson lab catalogue number, MPD strain ID, and where these strains were sequenced.

| Strain name | MGI strain ID | Jackson catalogue | MPD strain ID | Source |
| --- | --- | --- | --- | --- |
| C57BL/6J | MGI:3028467 | 664 | 7 | Reference |
| B10.D2-H2d/n2SnJ | MGI:2161770 | 462 | N/A | Stanford |
| BPL/1J | MGI:2164247 | 3006 | 19 | Stanford |
| BPN/3J | MGI:2162661 | 3004 | 27 | Stanford |
| CE/J | MGI:2159757 | 657 | 25 | Stanford |
| MA/MyJ | MGI:2159846 | 677 | 20 | Stanford |
| MRL/MpJ | MGI:2160037 | 486 | 58 | Stanford |
| NON/ShiLtJ | MGI:2163530 | 2423 | 34 | Stanford |
| NOR/LtJ | MGI:2162087 | 2050 | 106 | Stanford |
| NU/J | MGI:2161860 | 2019 | 127 | Stanford |
| P/J | MGI:2159762 | 679 | 55 | Stanford |
| PL/J | MGI:2159749 | 680 | 14 | Stanford |
| RBF/DnJ | MGI:2161383 | 726 | 33 | Stanford |
| RHJ/LeJ | MGI:2162860 | 1591 | 187 | Stanford |
| RIIS/J | MGI:2159809 | 683 | 26 | Stanford |
| SWR/J | MGI:2180845 | 689 | 12 | Stanford |
| TALLYHO/JngJ | MGI:3511696 | 5314 | 46 | Stanford |
| BTBR T<+> Itpr3<tf>/J | MGI:2162761 | 2282 | 1 | Stanford/Sanger |
| BUB/BnJ | MGI:2159907 | 653 | 24 | Stanford/Sanger |
| DBA/1J | MGI:2159759 | 670 | 65 | Stanford/Sanger |
| FVB/NJ | MGI:2163709 | 1800 | 9 | Stanford/Sanger |
| KK/HIJ | MGI:2161953 | 2106 | 18 | Stanford/Sanger |
| LG/J | MGI:2159748 | 675 | 78 | Stanford/Sanger |
| NZB/BINJ | MGI:2180844 | 684 | 11 | Stanford/Sanger |
| NZW/LacJ | MGI:2159914 | 1058 | 36 | Stanford/Sanger |
| RF/J | MGI:2159750 | 682 | 74 | Stanford/Sanger |
| SJL/J | MGI:2159739 | 686 | 17 | Stanford/Sanger |
| SM/J | MGI:2159787 | 687 | 31 | Stanford/Sanger |
| 129P2/OlaHsd | MGI:2164147 | N/A | 834 | Sanger |
| 129S1/SvImJ | MGI:3037980 | 2448 | 3 | Sanger |
| 129S5/SvEvBrd | MGI:3487126 | N/A | 62 | Sanger |
| A/J | MGI:2159747 | 646 | 4 | Sanger |
| AKR/J | MGI:2159745 | 648 | 15 | Sanger |
| BALB/cJ | MGI:2159737 | 651 | 5 | Sanger |
| C3H/HeJ | MGI:2159741 | 659 | 6 | Sanger |
| C57BL/10J | MGI:2159754 | 665 | 38 | Sanger |
| C57BL/6NJ | MGI:3056279 | 5304 | 182 | Sanger |
| C57BR/cdJ | MGI:2159792 | 667 | 32 | Sanger |
| C57L/J | MGI:2159746 | 668 | 37 | Sanger |
| C58/J | MGI:2159755 | 669 | 28 | Sanger |
| CBA/J | MGI:2159756 | 656 | 16 | Sanger |
| DBA/2J | MGI:2684695 | 671 | 8 | Sanger |
| I/LnJ | MGI:2159844 | 674 | 30 | Sanger |
| LP/J | MGI:2159761 | 676 | 10 | Sanger |
| NOD/ShiLtJ | MGI:2162056 | 1976 | 13 | Sanger |
| NZO/HILtJ | MGI:2173835 | 2105 | 122 | Sanger |
| SEA/GnJ | MGI:2159763 | 644 | 29 | Sanger |
| ST/bJ | MGI:2159751 | 688 | 69 | Sanger |

**Supplementary Figure 1.**
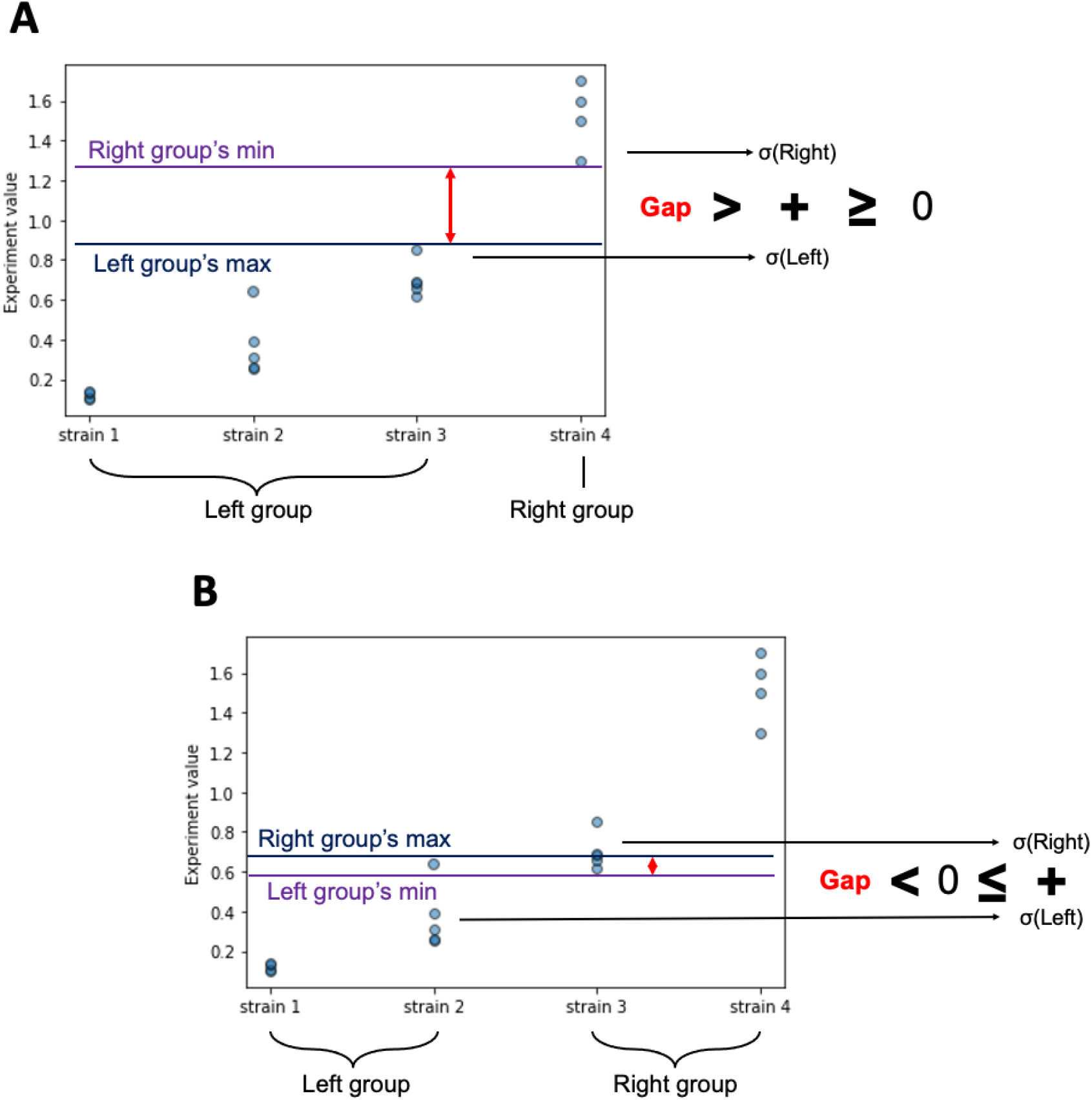
Phenotype separation. We first sort strains by average intra-strain phenotype value. Then we consider each boundary (n-1 if n is the number of phenotyped strains) to see if it is separable. To be separable, the right group’s minimum must be larger than the left group’s maximum (i.e., the gap is positive), and the sum of the standard deviations of the two strains flanking the boundary must be smaller than the gap size itself. **(A)** An example separable boundary between strain 1-3 and strain 4. **(B)** An example of a non-separable boundary.

**Supplementary Figure 2.**
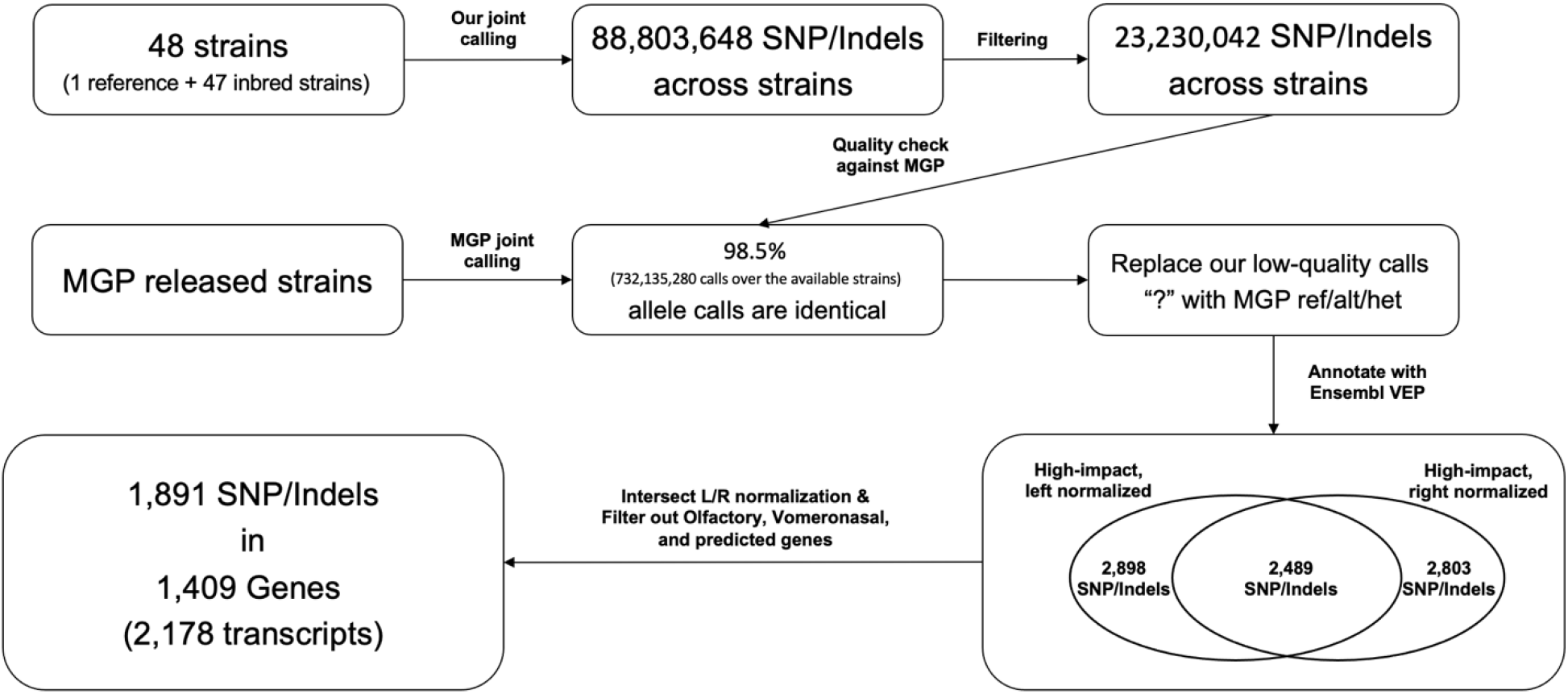
High-impact genomic variants. Using reference strain, C57BL/6J (MGI:000664), we called variants in 47 mouse strains (Supplementary Table 1). After filtering for quality, our calls were 98.5% identical to Mouse Genome Project (MGP) calls for strains that were available in MGP. We used their higher quality calls to augment ours, and then used Ensembl Variant Effect Predictor (VEP) to retain only high-impact variants such as stop gain or loss. Variants with more than one alignment were kept only if both alignments resulted in high-impact variants. Finally, we filtered out variants in large families of predicted and smell associated genes (see methods). The resulting set of high-quality candidate genes were matched against any phenotypic separation in Figure 1A.

**Supplementary Figure 3.**
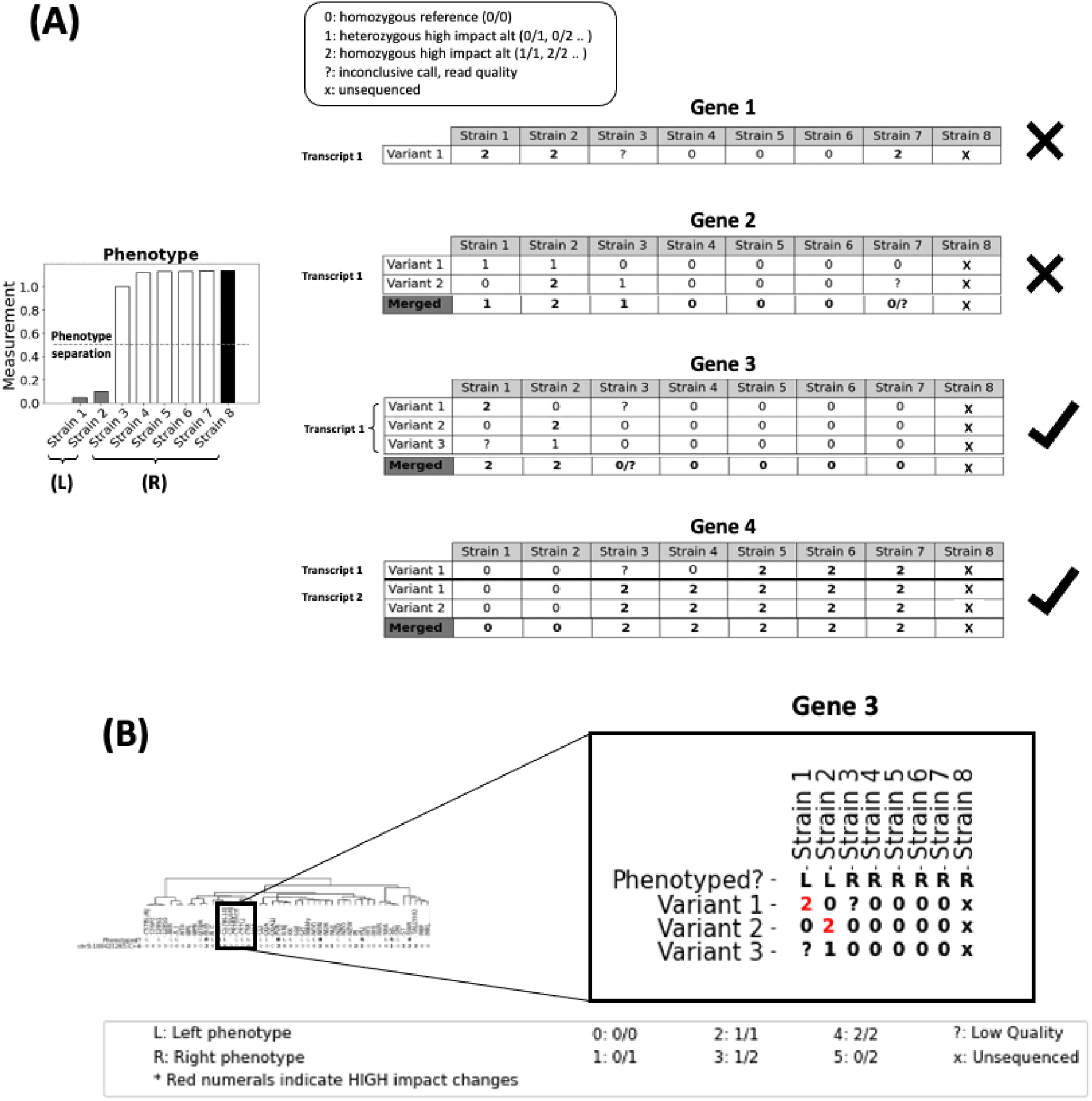
Phenotype-genotype matchings. **(A)** Left: An example experiment is found (automatically) to separate strains 1-2 from strains 3-8. Note that because the unit of measurement is arbitrary, calls of high/low, gain/loss are replaced simply by (L)eft and (R)right. Right: We tried to match four genes to the phenotype split. Gene 1 is a mismatch because of high-impact homozygous alternate calls in both groups (strains 1, 2, and 7). Gene 2 is a mismatch because of high-impact heterozygous alternate calls in both groups (strains 1 and 3). Gene 3 is a match (note that sequencing or genotyping strain 8 or a conclusive call in strain 3 can revoke the match). Gene 4 is also a match. **(B)** How gene 3 from (A) would be shown in our web interface (Figure S6).

**Supplementary Figure 4.**
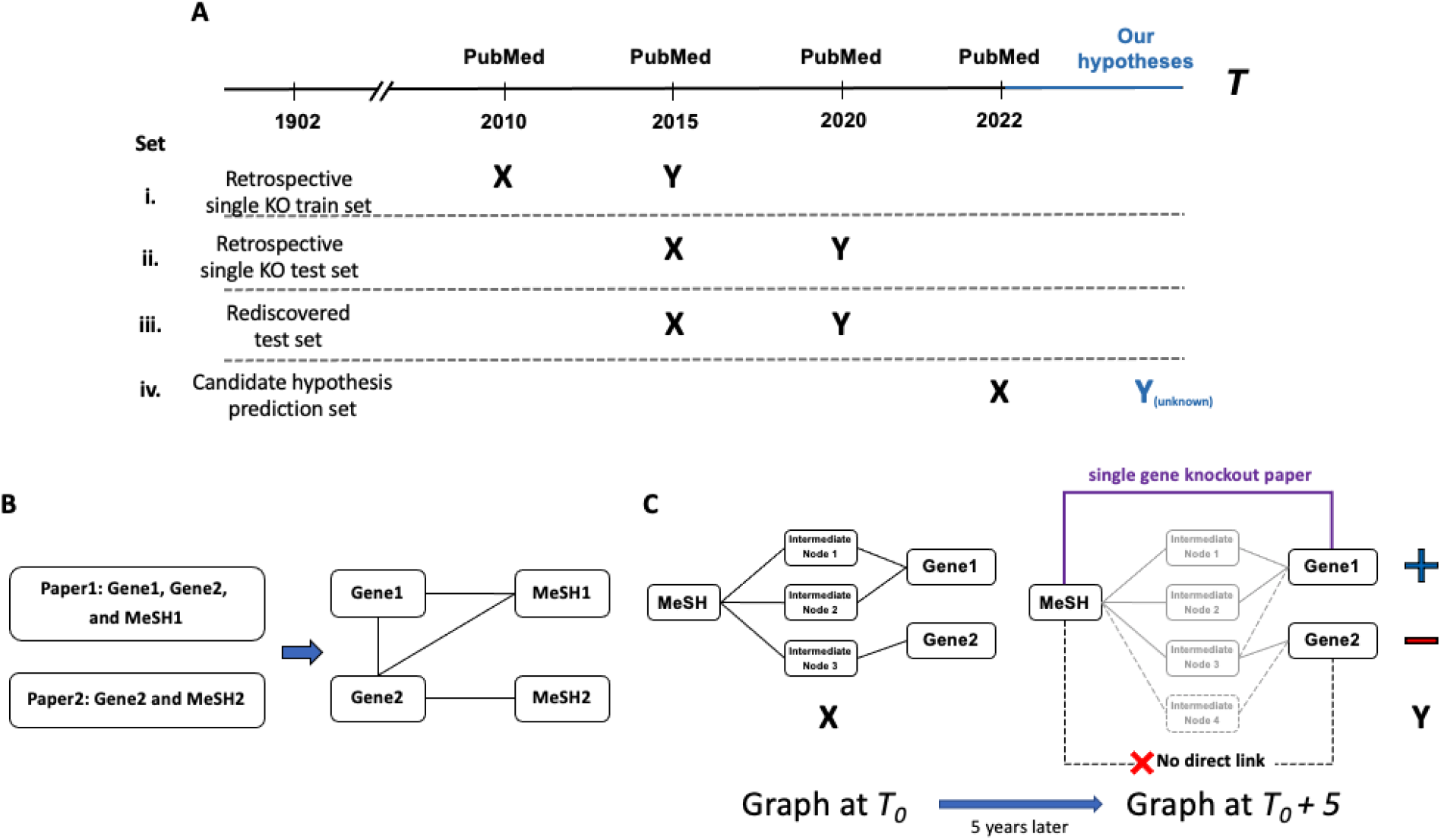
A machine learning framework to highlight our most promising novel testable hypotheses. **(A)** The classifier learns from examples in the retrospective single KO train set (i) to distinguish gene-phenotype relationships that became known in literature against gene-phenotype relationships that remained unknown (see methods). Performance of the classifier is measured on the retrospective single KO test set (ii) and our rediscovered test set (iii). Finally, the verified classifier is applied on our candidate hypothesis prediction set (iv) to prioritize the most promising candidates. **(B)** Each entity in the publication graph is either a MeSH term or Pubtator Central’s gene entity. For each paper, we obtained its MeSH tags, gene tags, and publication date. Undirected edges are extended between all MeSH terms and genes that are discussed in the same paper. **(C)** The retrospective sets are built using the open discovery framework on time-specific publication graphs (see methods). In the example, at time *T*_*0*_, the MeSH term shown is (only) indirectly linked to two candidate genes. By *T*_*0*_ *+ 5* years, only Gene1 was shown to be directly related (potentially causative) to the term. In the language of panel A, X = ((MeSH, Gene1), (MeSH, Gene2) and Y = ((+),(-)).

**Supplementary Figure 5.**
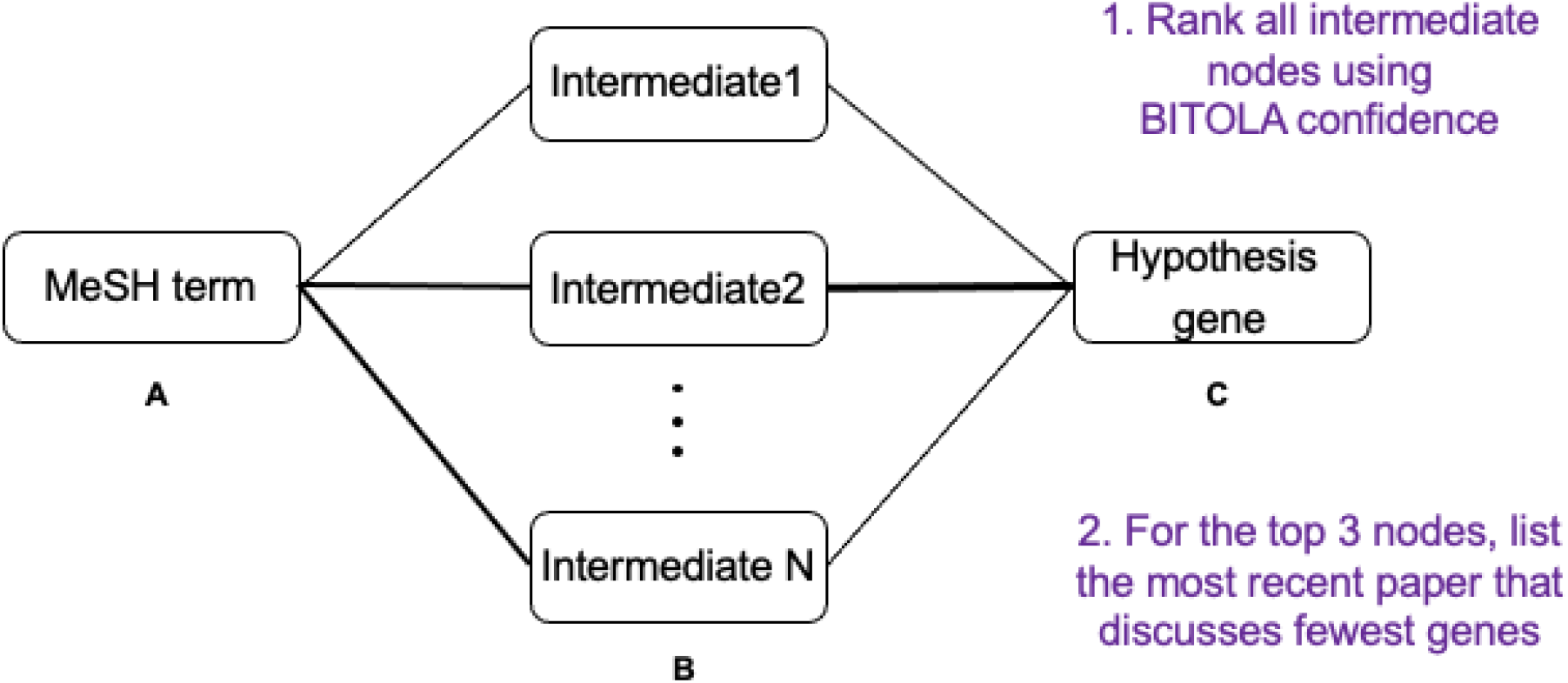
Explaining our classifier’s predictions using supporting literature. We ranked all intermediate nodes linking the phenotype term and the highlighted hypothesis gene using BITOLA confidence (see Methods). At the top would be intermediate terms most often co-mentioned with both phenotype and gene and therefore, possibly important in linking the two. For each evidence terms, our portal shows (Supplementary Figure 6) the most recent paper that mentions both term and gene.

**Supplementary Figure 6.**
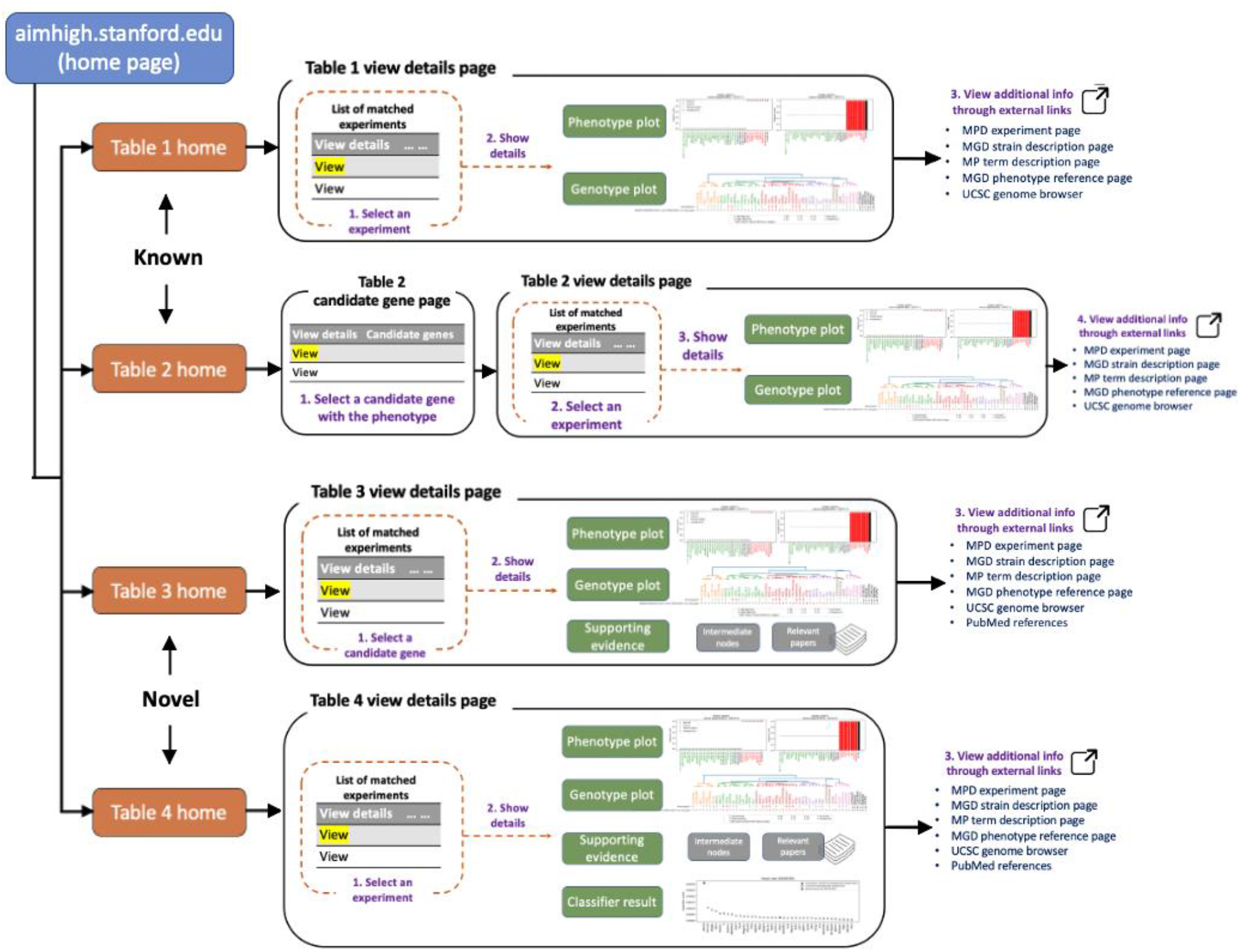
Roadmap of our web interface. We made all of our hypotheses available at aimhigh.stanford.edu. Each manuscript table (1-4) has a corresponding home page, formatted identically. The user can follow the purple instructions in each table to view details about each hypothesis, which include a phenotype plot with our derived boundary and a matching genotype plot. When applicable in Table 3 and Table 4, intermediate nodes and representative papers that may help evaluate the novel gene function hypothesis are also presented. In Table 4, our classifier result plot is also shown, with the selected gene hypothesis shown in red. Each page also has external links to the original data source (e.g., MPD and MGD) to aid in further evaluation.

